# *gilgamesh*, Drosophila casein kinase 1g, is required for myosin-dependent junction strengthening and epithelial folding

**DOI:** 10.1101/2024.09.11.612545

**Authors:** Lingkun Gu, Reina Koran, Jasneet Brar, Mo Weng

**Author notes:** equal contribution (Lingkun Gu, Reina Koran).

## Abstract

Adherens junctions, which serve as the primary physical link between cells, undergo remodeling in response to tension forces to maintain tissue integrity and promote tissue shape changes. However, the in vivo mechanisms driving this process remain poorly understood. Here, we identified Gilgamesh (Gish), the conserved fly homolog of casein kinase 1g as essential for myosin-dependent junction strengthening and tissue folding during apical constriction of Drosophila mesoderm. We show that Gish is recruited to spot adherens junctions in a contractile myosin-dependent manner. During apical constriction, Gish is required for junction strengthening by promoting growth and merging of small junction puncta, as well as stabilizing junction puncta at cell edges. The junction defects in Gish-depleted mesoderm result in breakage of the tissue-scale apical actomyosin network during apical constriction, and ultimately failure in mesoderm infolding. Our data show that Gish is a mechanosensitive kinase required for the integrity of adherens junctions during apical constriction.

## Introduction

Adherens junctions are the Cadherin-Catenin-based cell-cell adhesions that physically link the cytoskeletons of neighboring cells. Consistent with their intrinsic role in bearing mechanical tension, they are regulated by mechanical inputs in addition to biochemical signalings. The mechanosensitivity of adherens junctions enable them to be remodeled and adapt to the changing mechanical demand of the cell, which not only essential for keeping tissue integrity but also play a key role in cell shape changes (Campàs et al., 2023; Clarke and Martin, 2021; Guillot and Lecuit, 2013; Lecuit and Yap, 2015; Perez-Vale and Peifer, 2020; Pinheiro and Bellaïche, 2018). This is particularly important for tissue morphogenesis where cell and tissue shape changes happen extensively, most often driven by actomyosin contraction. However, the molecular mechanisms mediating the myosin-dependent remodeling of adherens junctions during tissue morphogenesis are largely unexplored.

Junction remodeling involves lateral clustering Cadherin-Catenin(Changede and Sheetz, 2017; Yap et al., 2015). The basic units of adherens junctions are lateral clusters of Cadherin-Catenin complexes with underlying actin network (Indra et al., 2018; Truong Quang et al., 2013; Wu et al., 2015). These nano-scale units further cluster to form a wide range of configurations, from puncta-like spot adherens junctions to belt-like zonula adherens. The major factor contributing to junction strength that can be regulated by the cell is the lateral clustering of Cadherin-Catenin complex. The number of Cadherins involved and the packing density differ greatly between different cell types and are remodeled when the tension status of the cell changes. The clustering of Cadherin-Catenin complexes depends on and also promotes the assembly of a special actin meshwork at junctions.

The adherens junctions have built-in mechanisms to sense and respond to tension: the core protein components, E-Cadherin and aCatenin, display confirmational changes when adherens junctions are under tension. The conformational change in the Cadherin extracellular domain enhances the trans-interaction between Cadherins in neighboring cells. The complex conformational changes of aCatenin recruits Vinculin, Ajuba, and Afadin/Canoe to adherens junctions, as well as directly increasing its interaction with actin filaments. Although several studies using cell culture demonstrate that vinculin is essential for junction integrity under tension (Huveneers and de Rooij, 2013; Le Duc et al., 2010; Leerberg et al., 2014; Thomas et al., 2013; Watabe-Uchida et al., 1998), such a role of vinculin appears to be minimal in intact animals such as Drosophila, Caenorhabditis elegans, zebrafish, or mouse (Alatortsev et al., 1997; Barstead and Waterston, 1991; Han et al., 2017; Maartens et al., 2016; Xu et al., 1998). Similarly, Ajuba does not appear to be required for embryogenesis in Drosophila, zebrafish or mouse suggesting a minor role in cell adhesions (Pratt et al., 2005; Razzell et al., 2018; Witzel et al., 2012), despite its prominent role in regulating tissue size(Rauskolb et al., 2014). The actin-binding protein Afadin/Canoe plays a significant role for adherens junctions during embryogenesis(Choi et al., 2013; Sawyer et al., 2009; Sawyer et al., 2011) but its mechanosensitivity appears to be limited to tricellular junctions(Sheppard et al., 2023; Yu and Zallen, 2020). These studies raise the question of whether the tension-dependent recruitment of junction interactors have functional significance or whether there are other such junction interactors yet to be identified.

We established previously that in apically constricting mesoderm epithelium of Drosophila embryos, spot adherens junctions are strengthened by growing in packing density and size in response to apical contractile myosin (Weng and Wieschaus, 2016). Adherens junctions serve as the essential anchors to connect the contractile actomyosin filaments in individual cells to a tissue-scale network that promotes the infolding of the mesoderm epithelium (Martin et al., 2010; Sawyer et al., 2009). The myosin-dependent strengthening is essential for junction integrity during the infolding because the polarity-based junction mechanism is downregulated in the mesoderm (Weng and Wieschaus, 2016). Both Vinculin and Ajuba are recruited to adherens junctions but as discussed before, none are required for embryogenesis.

To identify other potential mechanosensitive junction interactors, we looked for proteins that are recruited to adherens junctions during mesoderm apical constriction and are required for mesoderm infolding. We find that Gilgamesh (Gish), the fly homolog of casein kinase 1g, fulfills these two requirements. Gish and its homologs have been shown to be involved in various signaling pathways but they have never been implicated to be mechanosensitive or interacting with junctions(Chen et al., 2017; Gault et al., 2012; Li et al., 2020; Tan et al., 2010). We show that Gish recruitment depends on both adherens junctions and myosin contraction. Loss of Gish disrupts myosin-dependent junction strengthening and destabilizes the attachment of junction puncta to the cell cortex. As a result, apical constriction could not occur efficiently and mesoderm fails to fold to embryo interior.

## Results

### Loss of Gish leads to failure in ventral furrow formation

A supracellular network of contractile actomyosin is essential for the apical constriction and subsequent folding of Drosophila mesoderm epithelium during gastrulation(Heer et al., 2017; Martin et al., 2009). Such a tissue-wide network is formed by connecting actomyosin in individual cells through adherens junctions (Martin et al., 2010). To connect actomyosin filaments between cells without being pulled apart, adherens junctions are strengthened in response to the contractile myosin (Weng and Wieschaus, 2016) but the underlying molecular mechanism is not clear. We hypothesize that the proteins mediating such a myosin-dependent junction strengthening are recruited to adherens junctions in a tension-dependent manner and are required for successful folding of the mesoderm. We found that the GFP fusion protein from a protein trap line (Buszczak et al.) (Nerusheva et al., 2009) associated with the *gilgamesh* (*gish*) gene shows recruitment to junction-like puncta during mesoderm apical constriction. The fusion protein can indeed be detected by both anti-GFP and anti-Gish antibodies so we refer to it as Gish::GFP (Fig S1A). Gish is the sole fly homolog of highly conserved casein kinase 1g (CK1 g). It shares with the three human CK1 g about 70% identity in full length, and 80% identity in the kinase domain. CK1 g homologs, including Gish, are distinct from other casein kinases in their membrane association through the palmitoylation site at the carboxy terminus. Although Gish and CK1 g homologs have been shown to regulate many cellular processes, they are not known to be mechanosensitive. To test whether Gish is required for apical constriction and folding of mesoderm epithelium, we depleted Gish using two independent RNAi lines. Knocking down Gish using either RNAi line shows severe defects in mesoderm folding (Fig 1A). In wild type embryos, the mesoderm, which is marked by the expression of transcription factor Snail, is folded to the interior of the embryo. This folding leaves a linear closure on the surface of the embryo. By contrast, in *gish* RNAi embryos, mesoderm fails to be internalized (Fig 1A, B). Normally, mesoderm reenters cell cycle after they are folded inside (Foe, 1989). But in *gish* RNAi embryos, failed internalization leaves mesodermal cells dividing on embryo surface as indicated by the loss of nuclear Snail staining upon mitotic breakdown of nuclear envelope. To confirm the phenotype, we generated *gish* mutant embryos using transient CRISPR technique. We used anti-Gish antibodies to identify the embryos without Gish expression and found that these embryos display similar defects in mesoderm folding (Fig. S1B). These data show that Gish is essential for the proper internalization of mesoderm.

**Figure 1.**
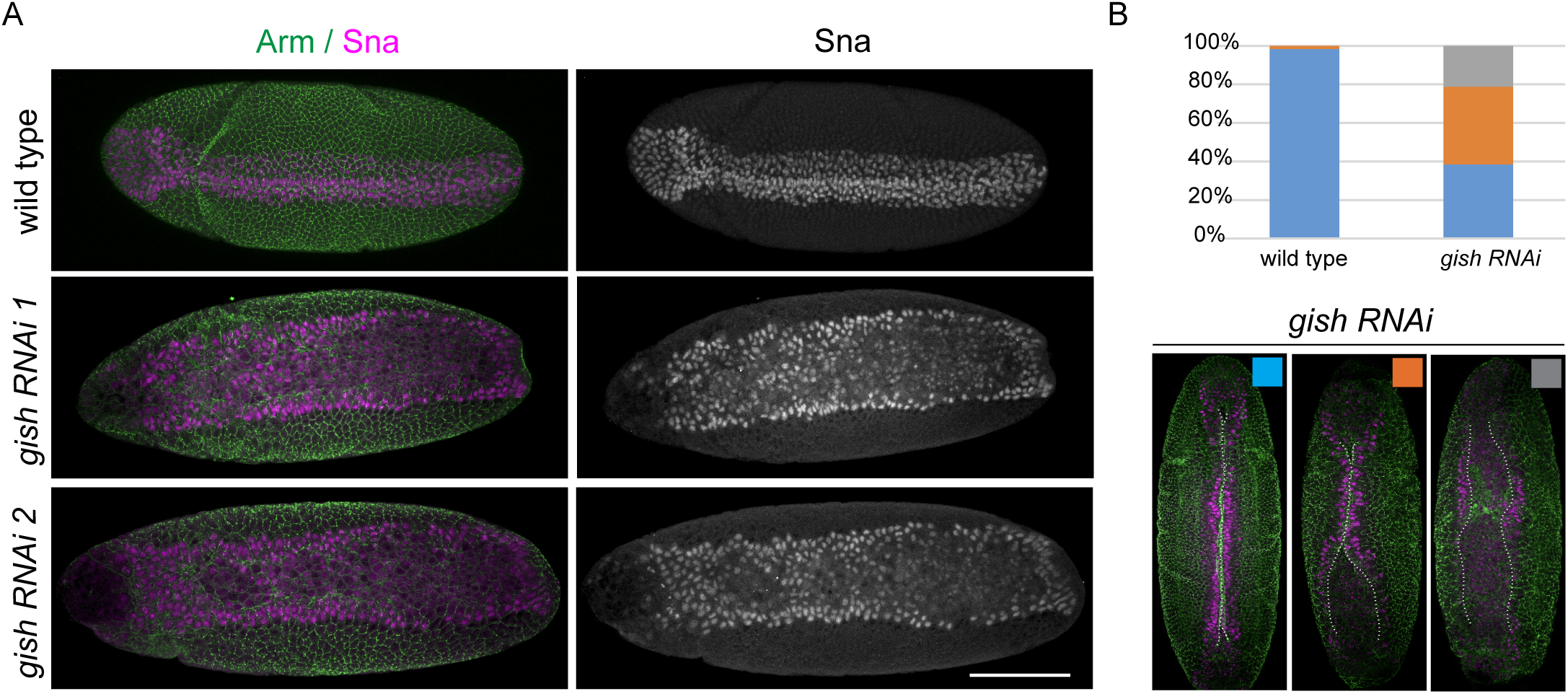
Gish is required for successful ventral furrow formation. (A) Wild type and gish RNAi embryos immunostained for Arm and Sna. Left: merged image with Arm shown in green and Sna in magenta; Right: Sna marks the mesodermal zone. Scale bar: 100 µm. (B) Quantification of mesoderm invagination defects in wild type and gish RNAi embryos. Embryos were categorized into three color-coded groups: Blue, >90% midline closure; Orange, partial midline closure; Gray, open. Wild type: n=62; gish RNAi n=52 and examples are shown below. White dotted line indicates the boundaries of the ventral furrow closure.

### Gish is recruited to adherens junctions-like puncta upon myosin contraction

We used Gish::GFP to characterize the junction recruitment of Gish protein during mesoderm apical constriction. Unlike the continuous belt-like adherens junctions that outline the cell, adherens junctions in Drosophila early embryos can be distinguished from cell-cell boundaries because they are in the form of light microscopy resolvable puncta which are tight clusters of E-Cadherin-Catenin complexes (Truong Quang et al., 2013). Shortly before gastrulation, such adherens junction puncta become smaller in the mesoderm due to loss of junctional polarity protein Bazooka/Par3 as part of the epithelial-mesenchymal transition process (Weng and Wieschaus, 2016) (Weng and Wieschaus, 2017). Upon apical constriction, adherens junctions are strengthened in a myosin-dependent manner: the junction puncta grow in size and brightness (Weng and Wieschaus, 2016) (Fig. 2A-A’, Movie 1). Gish, on the other hand, localizes uniformly to the cell membrane with little punctate localization before myosin is activated, consistent with being a membrane-associated protein (Fig. 2B-B’, left). This suggests that Gish does not localize to adherens junctions without myosin activation. However, as myosin is activated and mesoderm cells undergo apical constriction, Gish::GFP increasingly localizes to bright punctate structures, and the morphology and size of these Gish puncta resemble those of adherens junctions (Fig. 2B-B’, Movie 2). We used max projection to demonstrate all puncta within 7 um from apical surface while single z slice shows sharper membrane and punctate localization. We observed that the cells that activate myosin earlier acquire Gish puncta earlier, and as apical constriction progresses, all mesoderm cells acquire Gish puncta (Fig. 2B, right). Importantly, such puncta formation during apical constriction is not a general property of membrane proteins because other membrane markers P4M::mCherry and Resille::GFP remain uniformly membrane throughout apical constriction (Fig. 2C, Fig S2, Movie 3). This shows that puncta formation is not simply a result of shrinkage of cell edges or microscopic folds of cell membrane. E-Cadherin, Gish, and P4M all localize to plasma membrane but differ in the punctate localization during apical constriction (Fig. 2D). Overall, these data suggest that Gish displays distinct recruitment to adherens junction-like puncta when apical actomyosin contraction is activated.

**Figure 2.**
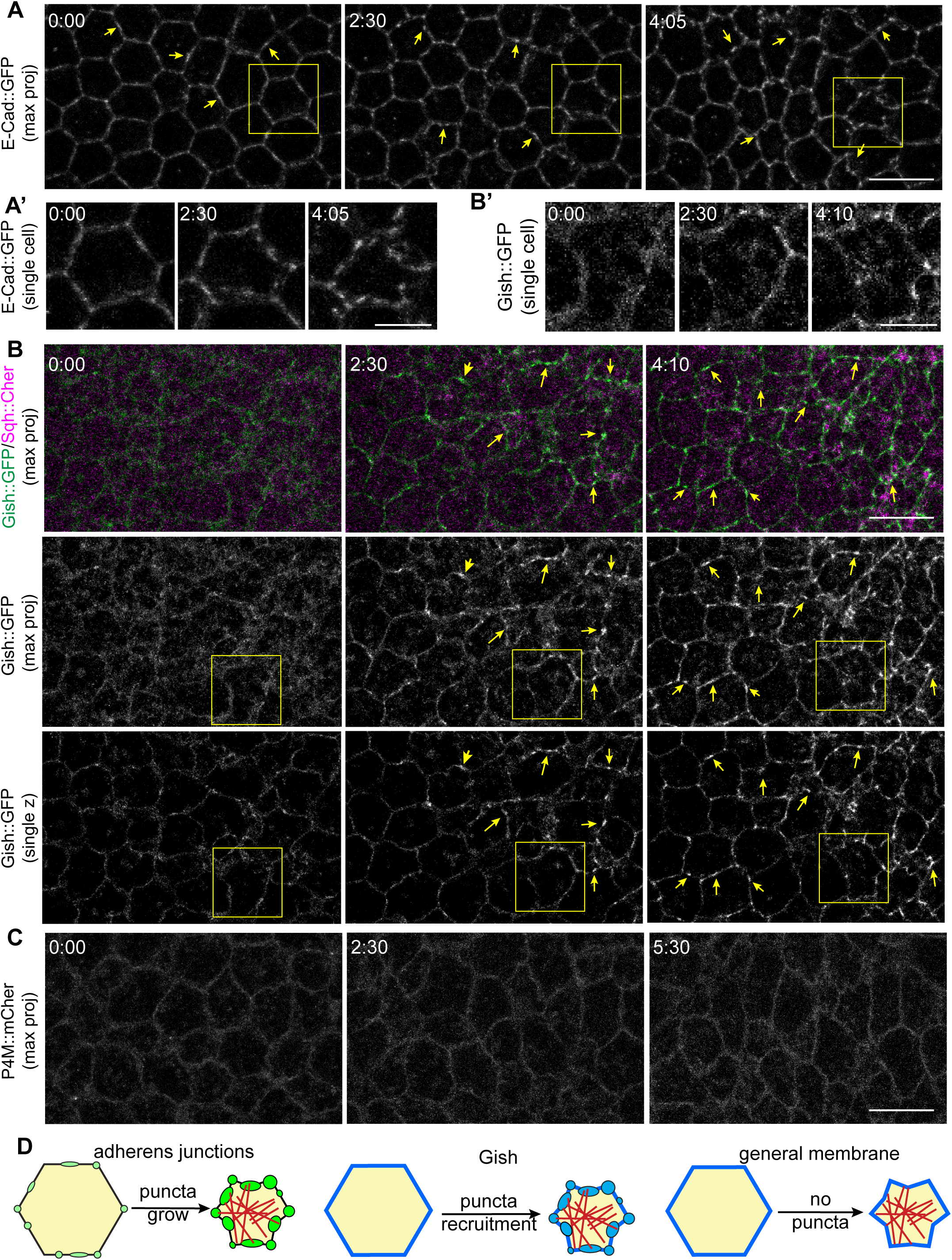
Gish is enriched into clusters subapically and apically during mesoderm apical constriction. (A-A’) Still images from time-lapse movies of ECad::GFP in the mesoderm. Arrows indicate adherens junctions puncta at the initial time point through the last time point, while the yellow boxes show an enlarged cell at the same time points. (A’): Enlarged images of the yellow boxed cell in (A). (B-B’) Still images from time-lapse movies of Gish::GFP (green) and Sqh::Cherry (magenta) in the mesoderm. (B) Top row: Max projection of Gish::GFP and sqh::Cherry merged images. Middle row: Max projection of Gish::GFP single channel images. Bottom row: single optical sections from confocal stacks of Gish::GFP. Yellow arrows indicate the same adherens junctions puncta at initial and later time points. Yellow boxes indicate the same cell in early and later time points. (B’) Enlarged images of the yellow boxed cell in (B). (C) Still images from time-lapse movies of P4M::mCherry in mesoderm. (D) Diagram illustrating the dynamic changes of the adherens junctions, Gish and P4M general membrane proteins in response to contractile myosin(red line). Scale bar for (A), (B), and (C): 10 µm; Scale bar for (A’) and (B’): 5 µm

### Gish puncta recruitment requires adherens junctions

To understand Gish puncta formation, we first determined whether such puncta localization of Gish requires adherens junctions. We were not able to compare junctions and Gish localization directly because Gish puncta are not preserved during fixation and live imaging markers of different colors for junctions and Gish are not available. However, we observe that Gish puncta resemble adherens junction puncta not only in morphology (Fig. 2B) but also in their behaviors. First, similar to adherens junctions, Gish puncta shift from subapical position to apical position during the early phase of apical constriction (Fig. 3A). Such shift of adherens junctions in apicobasal position is part of the myosin-dependent junction strengthening (Weng and Wieschaus, 2016), suggesting Gish is recruited to junctions at the onset of the strengthening process. Second, Gish puncta show similar anchor behavior for actomyosin filaments. During apical constriction, adherens junction puncta are pulled by actomyosin filaments towards the medial center of apical cortex, slightly bending the cell edges (Fig. 3B). Gish puncta show the same behavior of being pulled by the contractile actomyosin filaments (Fig. 3C). This strongly suggests Gish forms puncta through association with adherens junction structures.

**Figure 3.**
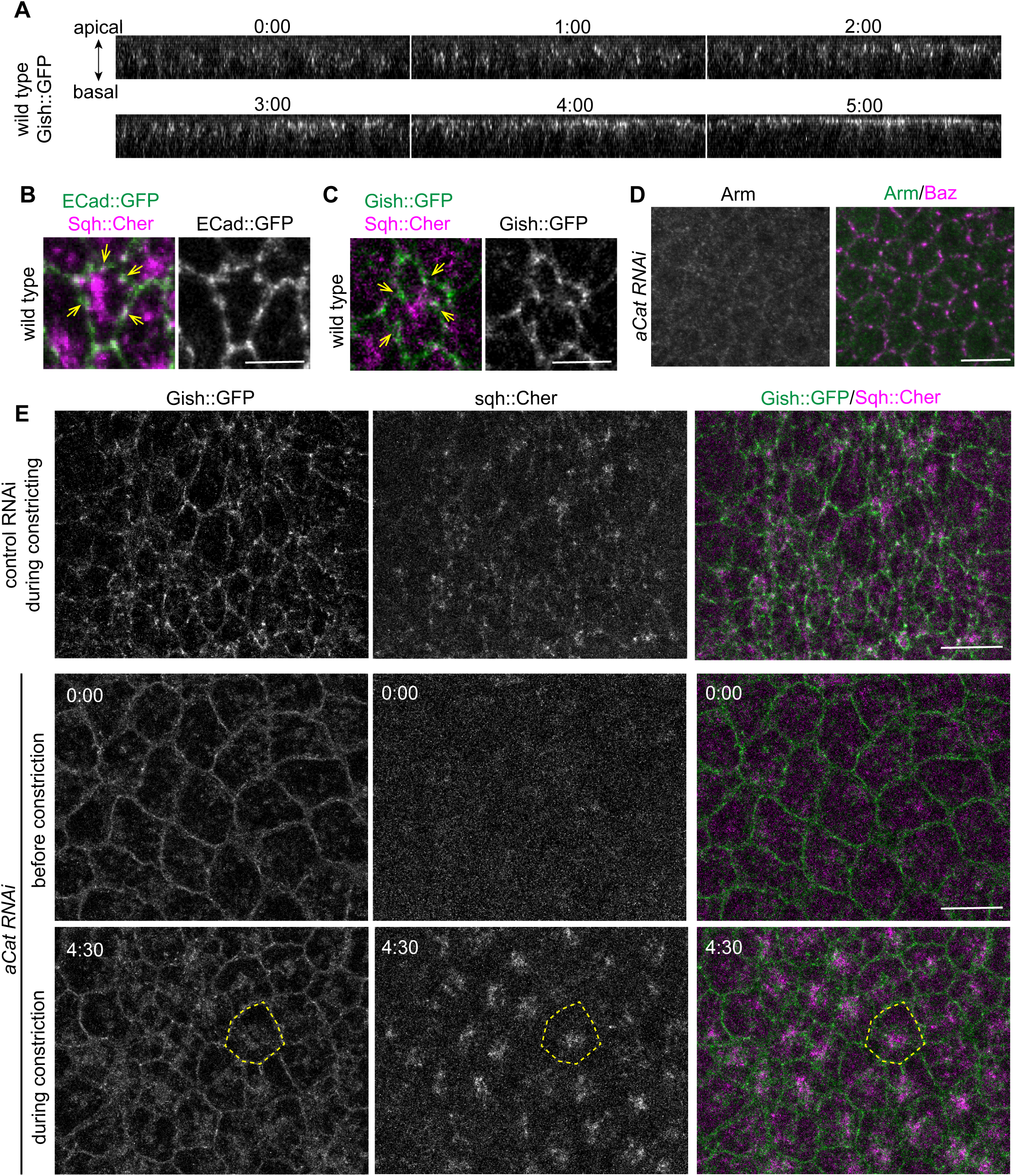
Loss of adherens junctions leads to mis-localization of enriched Gish puncta during apical constriction. (A) Still images (transverse view (YZ)) from time-lapse movies of Gish::GFP in the mesoderm during early apical constriction. Cross-section images are transverse views (YZ) generated from confocal stacks (XYZ), using the Reslice function in Fiji, showing the movement of Gish clusters along the apical-basal direction. (B) Still images from time-lapse movies of ECad::GFP and sqh::Cherry in the mesoderm during apical constriction. Yellow arrows: individual ECad puncta. (C) Still images from time-lapse movies of Gish::GFP and sqh::Cherry in the mesoderm during apical constriction. Yellow arrow: individual Gish puncta. (D) a-Catenin RNAi embryos immunostained for Arm (green) and Baz (magenta). Single Arm staining image shown on the left and the merged image shown on the right. (E) Still images from time-lapse movies of Gish::GFP and sqh::Cherry in control RNAi (top row) and a-Catenin RNAi (middle and bottom row) mesoderm. Single channel images show Gish::GFP (left) and sqh::Cherry (middle), with the merged image on the right. Dashed yellow lines: cell outline. Scale bar for (B) and (C): 5 µm; Scale bar for (D) and (E): 10 µm

Next, we tested whether Gish punctate formation depends on adherens junctions. When one of the core adherens junction components such as a-Catenin is knocked down by RNAi, the entire adherens junction structure is abolished (Fig. 3D). In a-Catenin RNAi embryos, Gish still uniformly localizes to the cell membrane but can no longer be recruited to puncta structures even as myosin shows strong accumulation on apical cortex (Fig. 3E, Movie 4). It has been shown previously that without adherens junctions, contractile actomyosin accumulates cell membrane into foci at the medial center of cell apical surface (Martin et al., 2010). Consistently, Gish, marking cell membrane, displays accumulation overlapping with myosin foci on apical surface (Fig. 3E, lowest panel, dotted line). Despite such foci-like accumulation of membrane Gish, no discrete bright puncta of Gish can be observed. Taken together, we show that the bright punctate form of Gish requires adherens junction structures as the platform.

### Myosin activation is necessary and sufficient for Gish puncta formation

As we have shown, the formation of Gish puncta in mesoderm cells only occurs upon apical constriction, and the cells undergoing apical constriction early acquire Gish puncta early. To test whether there is a similar association between Gish puncta and myosin activities in non-mesoderm tissues, we examined Gish localization in dorsal and lateral ectoderm epithelium at similar developmental stages. In dorsal epithelium where myosin is not activated, Gish only localizes to the plasma membrane but does not form puncta (Fig. 4A). In lateral epithelium where myosin is planar polarized, Gish also shows planar polarized recruitment (Fig. 4B).

**Figure 4.**
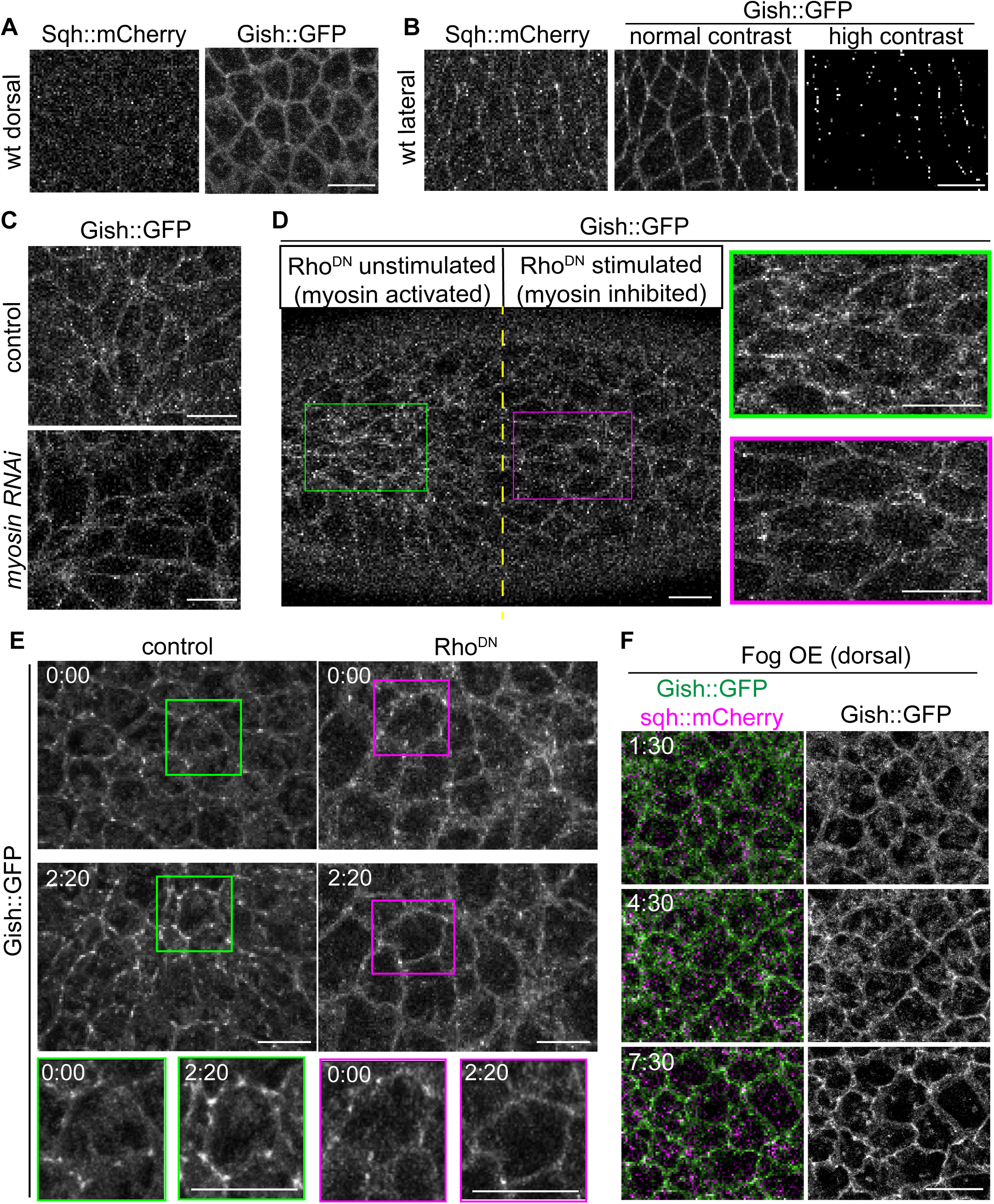
Myosin activation is essential for the enrichment of Gish at adherens junctions during gastrulation. (A) Still images from time-lapse movies of Gish::GFP and sqh::mCherry in dorsal ectoderm during ventral furrow formation. (B) Still images from time-lapse movies of Gish::GFP and sqh::mCherry in lateral ectoderm during germ band extension stage. The high contrast max projection image on the right shows the polarized Gish protein upon myosin activation. (C) Still images from time-lapse movies of Gish::GFP in control (top panel) and myosin RNAi (bottom panel) mesoderm during apical constriction. (D) Optogenetic inhibition of myosin activation by Opto-Rho1DN at beginning of Myosin activation in the mesoderm. Left of the dashed yellow line: no stimulation area of the embryo; Right of the dashed yellow line: stimulation area of same embryo by intermittent 488 laser light. Enlarged images of the green and magenta boxed regions are shown on the right. (E) Still images from time-lapse movies of Gish::GFP in the control (left panel) and optogenetically manipulated mesoderm during apical constriction. In the RhoDN images on right, myosin activation was constantly optogenetically inhibited by Opto-Rho1DN. Enlarged images of the boxed regions are shown at bottom. (F) Still images from time-lapse movies of Gish::GFP and sqh::mCherry in the dorsal ectoderm of Fog over expression embryos. Merged images shown on the left and single Gish::GFP channel images shown on the right. (A-F) Scale bar: 10 µm

These results imply that junctional recruitment of Gish depends on myosin activity. To test this hypothesis, we first knocked down myosin light chain Sqh using RNAi. In sqh RNAi embryos, Gish uniformly localizes to plasma membrane but could not form punctate structures at a stage where wild type embryos exhibit abundant Gish puncta (Fig. 4C). Due to its role in earlier stage development, myosin RNAi leads to deformed cells. To precisely control the timing of myosin inhibition, we used optogenetically controlled dominant-negative Rho (optoRho^DN^) (Guo et al., 2022). Stimulating optoRho^DN^ by 488 laser effectively inhibits myosin activation. First, we inhibited myosin shortly before apical constriction in one half of the imaging field. When the embryo develops to apical constriction stage, we can observe numerous intense Gish puncta along the cell peripheries on the side where myosin has not been inhibited (Fig. 4D, green). However, in the same embryo, much fewer Gish puncta can be detected on the side where myosin has been inhibited (Fig. 4D, magenta). This result shows myosin is required for the initial recruitment of Gish into puncta. Next, we tested if myosin is needed to maintain the Gish puncta. We inhibited myosin after some Gish puncta had already formed. In the control embryo, Gish puncta grow bigger and more intense as apical constriction progresses (Fig. 4E, green, Movie 5). By contrast, in the embryo where myosin is inhibited, already formed Gish puncta gradually become smaller and eventually disappear (Fig. 4E, magenta Movie 6). This suggests that the puncta form of Gish needs continuous myosin activity for maintenance.

Next, we tested whether activation of contractile myosin is sufficient to recruit Gish to adherens junctions. Ectopic expression of Fog is sufficient to induce myosin contraction through recruiting RhoGEF to the apical cortex in non-mesoderm tissues (Costa et al., 1994). We ectopically expressed Fog in the entire embryo. As myosin accumulates on the apical cortex, we observe recruitment of Gish into puncta in the dorsal epithelium (Fig. 4F) which normally does not show punctate Gish (Fig. 4A). Such global activation of myosin could not result in apical constriction since all cells pull on their neighbors, further showing that Gish puncta recruitment depends not on apical constriction but myosin activation.

### Gish is essential for the integrity of adherens junctions and supracellular actomyosin network

The recruitment of Gish to adherens junctions upon the onset of apical constriction and the dependency of such recruitment on myosin activation imply that Gish may play a role in the myosin-dependent junction strengthening. This hypothesis predicts that loss of Gish would lead to defects in adherens junctions specifically in mesoderm where the maintenance and strengthening of adherens junctions predominantly rely on myosin activity, but not in tissues such as the ectoderm where myosin is not activated and junctions rely on other mechanisms (Weng and Wieschaus, 2016). During apical constriction, the initially small junction puncta increase in size and packing density. Therefore, disruption of myosin-dependent junction mechanism should lead to smaller or loss of adherens junction puncta in the mesoderm. In addition, since adherens junctions are the essential anchors for tissue-wide actomyosin network, weakened junctions would compromise the integrity of the actomyosin network (Martin et al., 2010).

To assess the state of adherens junctions in *gish* RNAi embryos, we first examined endogenous adherens junctions in highly constricted cells using anti-Armadillo (Arm, fly b-Catenin) antibody. In wild type apically constricted mesoderm, the strengthened adherens junctions appear as bright and large puncta, elongated covering most of the cell edges (Fig. 5A, Arm, Arm/cell outline). These junctions connect actomyosin into a continuous supracellular network (Fig. 5A, Arm/Zip(Myo)) which bends the epithelium (Fig. 5A, far right). In striking contrast, adherens junctions in *gish* RNAi mesoderm appear as numerous small puncta loosely gathered together into foci, mostly localizing on the apical surface rather than the cell edges (Fig. 5B, Arm, Arm/cell outline). We used the dim membrane signal of Arm 1 um below the junction region to label the cell outline and found that the foci and the gaps around them can involve multiple cells. Consistent with the altered adherens junctions which serve as actomyosin anchors, actomyosin filaments no longer form a continuous network across cells but coalesce into disconnected foci together with the small junction puncta (Fig. 5B, Arm/Zip). Such breakage of supracellular myosin network results in ineffective tissue bending (Fig. 5B, far right). More and larger gaps of myosin network across multiple cells appear in embryos at later stages of apical constriction while the mesoderm epithelium only bend minimally (Fig. 5B, lower panel). The abundance of small junction puncta suggests Gish may not be required for the initial formation of small junction puncta, although we cannot rule out that this is due to an incomplete depletion of Gish. Importantly, these phenotypes are specific to apically constricting mesoderm: adherens junctions still localize to the cell edges as bright puncta in flanking ectoderm despite the severe defects in mesoderm of the same embryo (Fig. S3A). This indicates the apical surface gathering of junction puncta is not a general consequence of loss of *gish* but a defect in cells where myosin-dependent mechanism is essential for intact adherens junctions.

**Figure 5.**
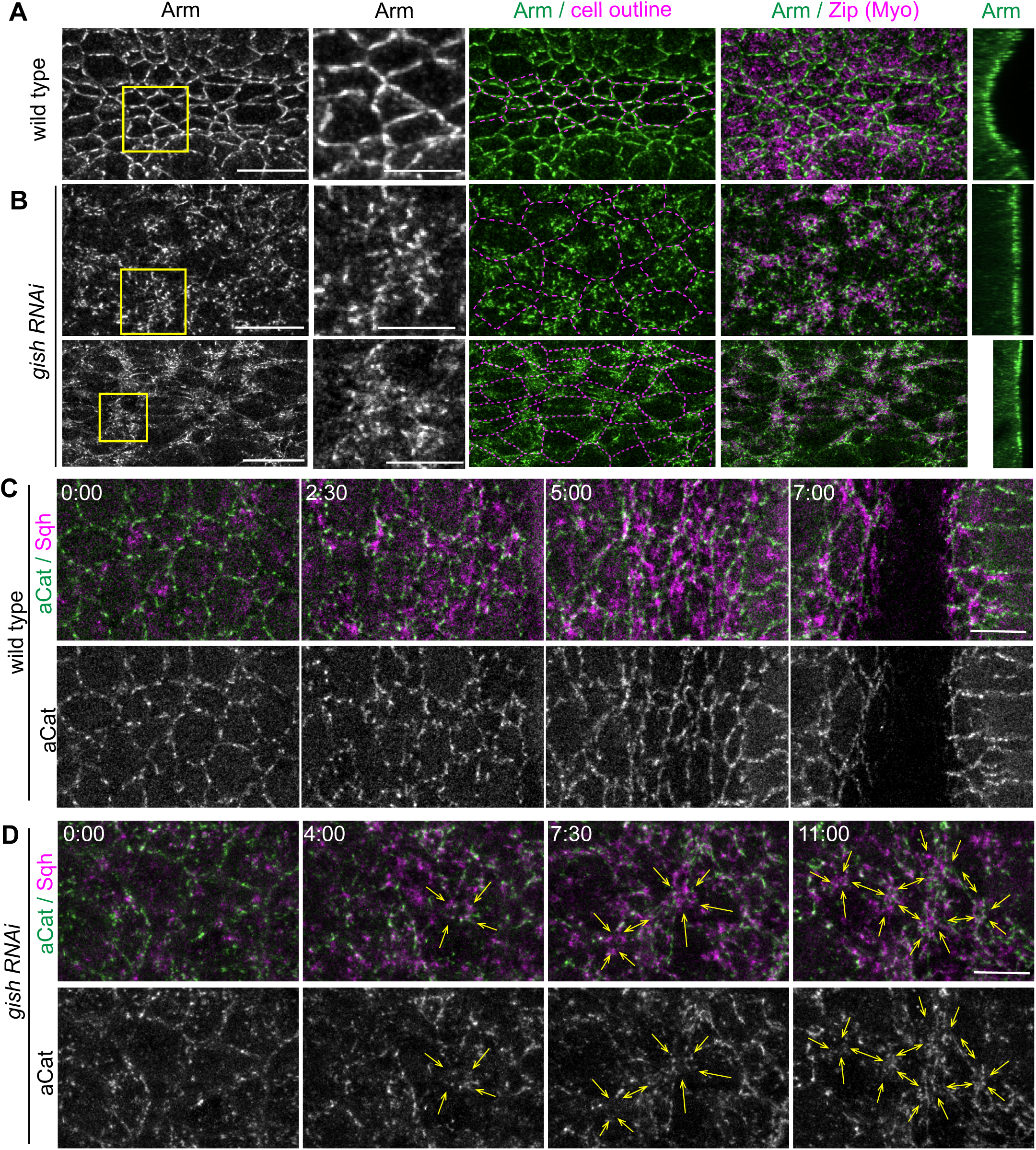
Loss of Gish disrupted the supracellular actomyosin network. (A), (B) wild type and gish RNAi embryos immunostained for Arm and Zip respectively. The enlarged yellow boxed region on the leftmost column is shown in the second column. The artificial dotted magenta cell outline is based on subapical Arm signal shown at middle column. Cross section view (YZ) showing the depth of ventral furrow in the last column. Anterior to posterior orientation is shown horizontally. Scale bar is 10 µm except the enlarged image is 5 µm. (C), (D) Still images from time-lapse movies of aCat::Venus and sqh::mCherry in the WT and gish RNAi mesoderm. Yellow arrows: junction foci formation in gish RNAi mesoderm. Anterior to posterior orientation is shown vertically. Scale bar: 10 µm

To follow how the junction defects and breakage of myosin-junction network are developed over time, we imaged adherens junctions in living embryos using aCatenin (aCat)::Venus since we find E-Cad::GFP to be incompatible with *gish* RNAi. In wild type embryos, junction puncta are constantly pulled by contractile actomyosin but they do not deviate significantly from the cell outline throughout apical constriction (Fig. 5C, Fig. S3B, Movie 7). In *gish* RNAi embryos, more and more junction puncta show localization to the apical center of the cell as apical constriction progresses (Fig. 5D, Fig. S3B, Movie 8). These junction puncta from different cell edges appear to be gathered into foci by the contractile myosin but also separate as the myosin-junction foci undergo constant breakage and reconnection. As myosin contraction increases, more foci and larger gaps are developed. Consistent with the observation from fixed embryos, junction puncta are abundant at later stages of apical constriction, but they remain small in size.

Similar to embryos with junctions depleted (Martin et al., 2010)(Fig. 3E), these myosin foci in *gish* RNAi embryos accumulate membrane blebs and tethers as shown by resille::GFP and scanning EM (Fig. S3C-D). Such a phenomenon in junction-defective embryos is proposed to be due to residual junctions pulled by contractile myosin in one cell while detached from the actin cortex from the neighboring cell. The similarity with junction-depleted embryos in membrane abnormality implies that junctions are weakened in *gish* mutant mesoderm.

These data show that loss of gish results in defective organization of adherens junctions, which in turn leads to breakage of the tissue-wide myosin-junction network.

### Gish is required for junction puncta to grow and merge

Our previous study shows that adherens junctions in mesoderm are strengthened in response to contractile myosin through incorporating more Cadherin-Catenin complexes into junction puncta and merging of small puncta (Weng and Wieschaus, 2016). Based on the above observation, we hypothesize that the abundance of small puncta in *gish* mutant embryos and their abnormal displacement to apical center may be due to the failure of junction puncta to grow or merge. To follow the fate of individual junction puncta, we switched to Ajuba(Jub)::GFP to visualize adherens junctions because aCat::Venus does not allow high temporal resolution due to rapid photobleaching. Jub::GFP has been shown to label adherens junctions in a myosin-dependent manner (Rauskolb et al., 2014; Razzell et al., 2018). In wild type embryos, Jub::GFP-labeled adherens junctions display the same myosin-dependent strengthening behaviors during apical constriction. Initially, the puncta are small and scarcely distributed along cell edges. As apical constriction progresses, many more puncta accumulate on each cell edge. At late stages of apical constriction, adherens junctions appear less punctate and more continuous covering each cell edge (Fig. 6A, Movie 9). In *gish* RNAi embryos, the Jub-labeled junction puncta do arise and even accumulate to a high amount. However, throughout apical constriction, the cells only display small puncta despite them gathered close into foci, consistent with the observation using aCat::Venus (Fig. 6B, Movie 10). Such a phenotype of small puncta and loss of cell edge association can be observed in embryos with mild gish RNAi judged by the delayed but complete infolding of mesoderm (Fig. 6C, Movie 11).

**Figure 6.**
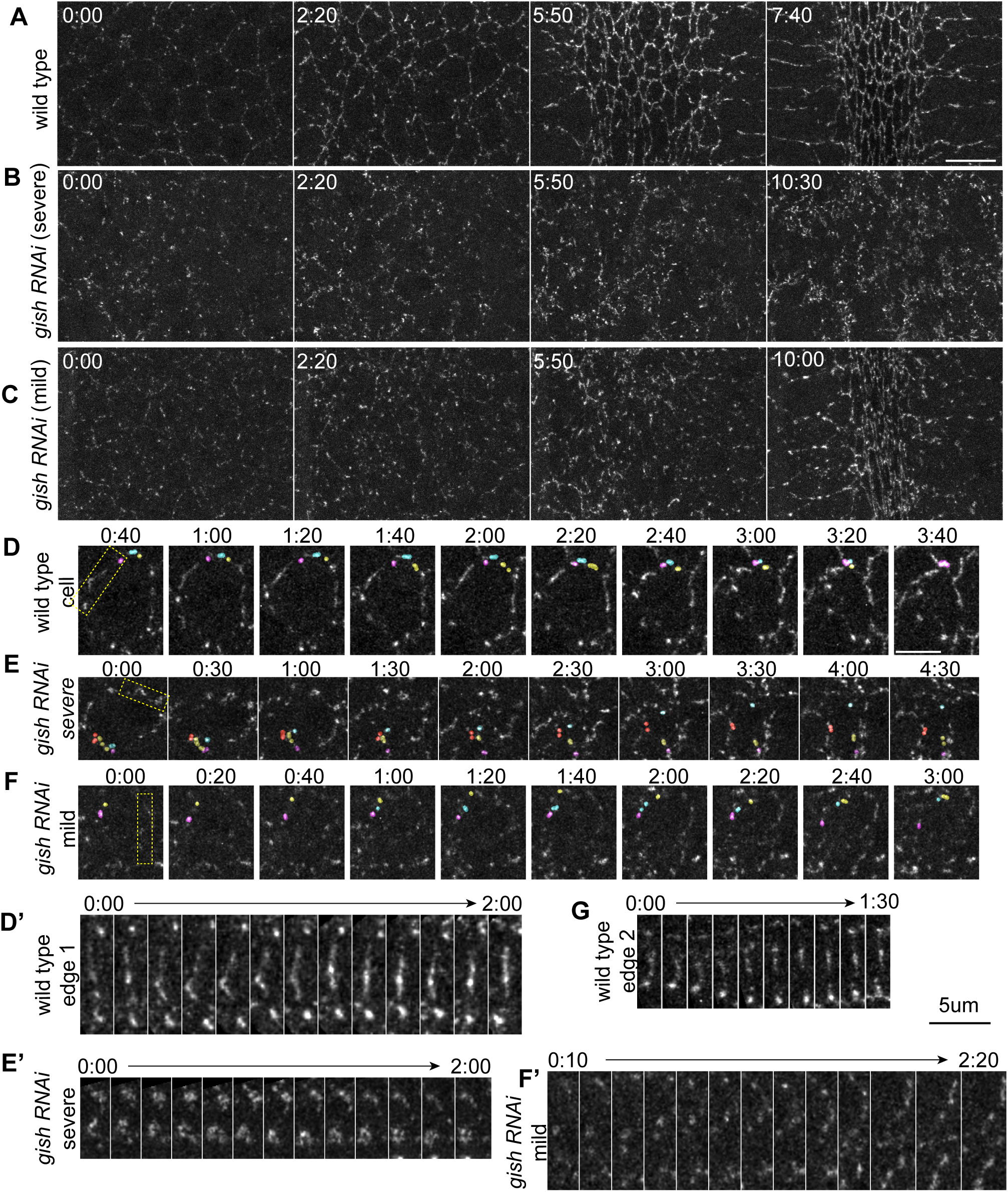
Gish is required for adherens junctions strengthening in mesoderm cells during apical constriction. (A), (B), and (C) Still images from time-lapse movies of Ajuba::GFP in the mesoderm of wild-type and gish RNAi embryos, respectively. Scale bar: 10 µm. (D), (E) and (F) Single cell still images from time-lapse movies of Ajuba::GFP in the mesoderm of wild-type and gish RNAi embryos, respectively. Different color coded Individual Ajuba clusters merged into a big cluster in WT but not in gish RNAi embryos. (D’-F’) Tracking of Ajuba clusters in an individual cell edge (yellow dashed box in D-F, respectively) during apical constriction. (G) Ajuba cluster grows in size and intensity even cell edges extend. Scale bar for (D-G): 5 µm.

We followed individual junction puncta to observe the growth and merging behavior of the puncta (Fig. 6D-G). In wild type cells, small puncta gradually arise, grow in size and those on the same edge merge into bigger puncta (Fig. 6D-D’, Movie 12). As a result, the appearance of cell edges change from a string of puncta to a more continuous line as apical constriction progresses. When an edge is pulled by contracting actomyosin, the string of puncta bends slightly but the puncta stay close together or merge (Fig. 6D’, the first half of the tracking period). Junction puncta can grow or merge even when cell edges expand, again indicating the growth of junction puncta is not simply due to concentrating junction materials as cell edges shrink (Fig. 6G). By contrast, Jub-labeled junction puncta in *gish* RNAi embryos do not show as much growing and merging and cell edges remain to be strings of puncta (Fig. 6E-F, E’-F’, Movie13-14). The string of puncta is often broken when the cell constricts, obscuring the cell outline and leading to accumulation of puncta at the apical center of the cell. The junction puncta originated from the same cell edge are increasingly pulled far apart from each other as apical constriction proceeds. Instead, junction puncta coming from different edges come together to form the loosely associated myosin-junction foci. Such significant deviation from the edge the puncta originally resided on suggests detachment of adherens junction puncta from the actin cortex and is consistent with the membrane tether and blebbing associated with the myosin-junction foci (Fig. S3C-D).

Taken together, we show that Gish is essential for the growth and merging of small junction puncta into bigger ones. Without Gish, junction puncta in apical constricting mesoderm could not form continuous line-like junctions along the cell edges and individual small puncta pulled by actomyosin contraction are displaced dramatically to the apical center of the cell. These junction defects lead to breakage of tissue-scale myosin network and failed tissue infolding.

## Discussion

We have demonstrated that the fly casein kinase 1g Gish is recruited to adherens junctions in response to myosin contraction and is essential for the mechanosensitive junction strengthening during the infolding of mesoderm epithelium. In Gish RNAi embryos, the myosin-dependent junction strengthening is disrupted. During the apical constriction of Gish RNAi mesoderm, small junction puncta accumulate to a high number but cannot grow or merge into bigger and more continuous adherens junctions along the cell edges. Instead, these small puncta are displaced to the center of the apical membrane forming foci with contractile actomyosin. This disrupts the formation of a continuous tissue-scale actomyosin network, leading to failed tissue folding.

The small junction puncta in gish RNAi embryos suggests Gish promotes junction puncta growth and merging. Junction puncta growth and merging are the result of further clustering of Cadherin-Catenin complexes. Since Cadherin clustering requires the assembly of actin network directly underneath the Cadherin clusters(Campàs et al., 2023), Gish may be recruited to promote actin assembly at the junction puncta. Alternatively, Gish recruitment may enhance binding of Cadherin-Catenin complexes to actin filaments and therefore indirectly increase junctional actin network for clustering.

Another junction defect in gish RNAi embryos is their apparent detachment from cell edges. By comparing wild type and gish RNAi embryos, it is clear that normally junction puncta stay closely associated with cell edges even when pulled by medial contractile actomyosin. Although cell edges in wild type embryos can be bent when junction puncta are pulled on one side, cell edges do not deform drastically and soon recover. This implies the existence of an actin structure that provides rigidity to the cell circumference. Indeed, Diaphanous, activated by Rho, has been shown to localize to and assemble actin cables around apical cell circumference during apical constriction of fly mesoderm(Mason et al., 2013). This could be a good candidate actin structure that supports cell edge rigidity. The dramatic displacement of junction puncta in *gish* RNAi embryos suggests these puncta are connected to the medial actomyosin but detached from the circumferential actin cables. So Gish could function to enhance the interaction between junctions and the circumferential actin cables.

The two junction defects – small size and detachment from cell circumference – could be independent phenotypes or one could be the consequence of the other. Detachment could disrupt merging since it pulls junction puncta away from other puncta. It is also possible for small size itself to lead to detachment. Junction detachment has been observed in embryos with depleted junctions(Martin et al., 2010). These embryos show similar though not identical myosin and membrane foci as well as membrane tethers on apical center of the constricting mesoderm cells. However it is not clear if the residual adhesion that leads to such myosin and membrane foci is residual adherens junctions. In addition, junction puncta in *gish* RNAi embryos are much bigger than the virtually undetectable cell adhesion in embryos with depleted junctions. It will be interesting to see whether embryos with moderately depleted but detectable junctions show similar junction detachment phenotype.

## Methods

### Genetics

To generate transient CRISPR mutant embryos: flies carrying gRNA against gish (BDSC 84029) were crossed to flies carrying nanos-Cas9 to induce mutant alleles in germ line (Kondo and Ueda, 2013)(Port et al., 2014). Females from such a cross were used to produce gish mutant embryos. Mutant embryos were identified by the lack of Gish protein using anti-gish antibody (Tan et al., 2010).

### Embryo fixation, immunostaining, and imaging

Drosophila embryos were collected on apple juice agar plates at 25°C for 2 h, followed by an additional 2-h aging period at the same temperature after the removal of adult flies. Then the embryos were dechorionated with 50% of 8% household bleach (4% sodium hypochlorite) and fixed by the heat-methanol protocol as described before (Müller and Wieschaus, 1996). Briefly, in a 15 ml glass vial, the dechorionated embryos were incubated in 3 ml of hot salt solution (0.4% NaCl, 0.03% Triton x-100) for 10 s, and then swiftly cooled by adding 2 volumes of prechilled salt solution. Discarded the salt solution from the vial and then added a 1:1 mixture of methanol and heptane. Vortexed vigorously for at least 30 s to remove the vitelline membranes. Then, the embryos were transferred into an Eppendorf tube washed with methanol three times. The embryos were stored in methanol at −20°C until use.

Embryos were incubated in 10% BSA blocking buffer for 1 h, followed by overnight staining with primary antibody at 4°C and subsequent staining with the secondary antibody for 2 h at room temperature. Following antibody staining, embryos were sorted and mounted in Aqua-PolyMount (Polysciences).

Images are acquired on a Zeiss LSM800 confocal microscope equipped with high-sensitive GaAsp detectors. LD LCI plan-Apochromat 25x/0.8 objective was used for whole embryo images and plan-Apochromat 63x/1.4 NA Oil DIC M27 objective was used for high magnification images.

### Live imaging and optogenetics

Embryos for live imaging were prepared using the described protocol (Gu and Weng, 2021). All experiments were performed at RT for live imaging. Images were acquired on a Zeiss LSM800 confocal microscope equipped with high-sensitive GaAsp detectors. LD LCI plan-Apochromat 25x/0.8 objective was used for low magnification images and plan-Apochromat 63x/1.4 NA Oil DIC M27 objective was used for high magnification images. Diode 488 and 561 nm lasers were used to excite GFP and mCherry, respectively. The pinhole was set at 1 Airy unit for 488 for all images. Zeiss Definite Focus was used to maintain the focal plane.

For the optogenetic experiment, the opto-Rho1DN optogenetic tool from Bing He’s lab (Dartmouth College) was used in this study. All fly crosses were kept in the dark. Live samples were prepared under red light. The embryo stages were determined by monitoring the myosin signal using a 561 nm laser. For Fig. 4D, just before noticeable actomyosin activity was observed, 0.1% 488 nm laser was continuously applied to stimulate the optogenetic module within defined ROI for 10 s. After a 50 s interval, the ROI was stimulated again. This process was repeated for a total of 5 cycles. At the end of the stimulation, post-stimulation images were acquired, including those of the no-stimulation region. For Fig. 4E, 1.5% 488 nm laser and 561 nm laser were constantly applied to the ROI with a temporal resolution of 10 s.

### SEM

Embryos were dechorionated with 50% of 8% household bleach (4% sodium hypochlorite) and fixed for 25 min with 25% glutaraldehyde in 0.1 M sodium cacodylic buffer and heptane. The vitelline membrane was then manually removed in PBS with a needle, and embryos were dehydrated by a gradient of ethanol concentration (25%, 50%, 75%, 95%, and 100%). Embryos were then incubated for 10 min in a 1:1 mixture of ethanol and hexamethyldisilazane (HMDS), followed by two additional incubations with 100% HMDS. After HMDS evaporated completely, embryos were transferred to the SEM stub and gold coated using a Sputter Coater 108auto (Cressington). Samples were imaged using Hitachi TM-1000.

### Imaging processing and analysis

All images for publication were processed in ImageJ (http://rsb.info.nih.gov/ij/). Brightness and contrast were adjusted for the whole image. Quantitative analysis was done using MATLAB (MathWorks).

### Quantification of embryo phenotype

The fixed embryos, stained with Arm and Sna, were manually categorized based on their phenotype.

For scoring ventral furrow closure phenotypes (Fig. 1B), embryos were sorted according to the percentage of the ventral furrow that was closed. Embryos with >90% fused/sealed ventral furrow were classified as having a complete closure. Embryos with <10% sealed ventral furrow were classified as open. The remaining embryos were classified as partial closure. Embryos from late stage 6 and stage 7 were used for quantification. Ventral furrows in wild type embryos usually close by late stage 6 and become less distinguishable by stage 8 when domain 14 enters the cell cycle.

### Western blot

Drosophila embryos were collected on apple juice agar plates at 25°C for 2 hours, followed by an additional 2-hour aging period at the same temperature after removing adult flies. Then the embryos were dechorionated by 50% of 8% household bleach (4% sodium hypochlorite). Embryo extraction was conducted by incubating dechorionated embryos in 2x Laemmli buffer (Bio-Rad) with 5% 2-mercaptoethanol at 95°C for 2 min. The lysates were then cleared by centrifugation at 16,000g for 5 min. 8% surePAGE gels (GenScript) were used for electrophoresis and Immobilon-FL PVDF membranes (Millipore) were used for transfer. Membranes were blocked for 1 h in PBST (0.1% Tween-20 in PBS) containing 10% dry milk. Then the membranes were incubated in primary antibody overnight at 4°C (Guinea pig anti-Baz, 1:500; Mouse anti-Tubulin, 1:2000) and subsequently in secondary antibody for 1 hour at room temperature (IRDye-coupled secondary antibody: Donkey anti-Gp 800CW Cat# 926-32411 and Goat anti-Mouse 680RD Cat# 926-68070, 1:5000). Membranes were visualized with an Odyssey infrared imaging system (LI-COR Bioscience), and brightness and contrast were adjusted for the whole image using ImageStudio software.

