## Supplementary material for "*gilgamesh*, Drosophila casein kinase 1g, is required for myosin-dependent junction strengthening and epithelial folding": supp. figures

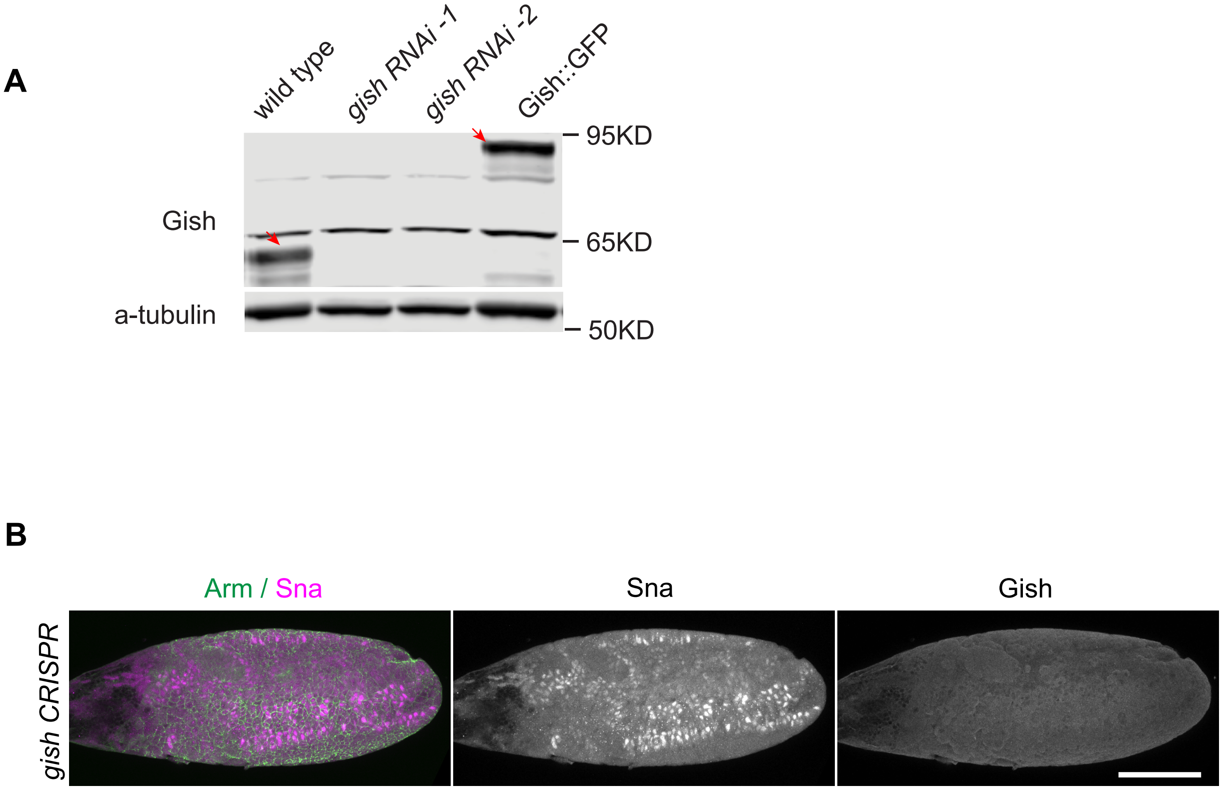


Figure S1.  Knocking out Gish in fly embryos leads to defects in gastrulation.

(A) Western blot of Gish protein in wild type, GFP-tagged Gish line, and gish RNAi embryos. ő-tubulin served as the loading control. Red arrows indicate endogenous Gish and GFP-tagged Gish protein in wild type and Gish::GFP embryos respectively. (B) gish CRISPR knockout embryos immunostained for Arm, Sna, and Gish. Left: Arm and Sna merged image; Middle: Sna staining to show mesoderm; Right: Gish staining showing Gish protein is depleted. Scale bar: 100 Õm.

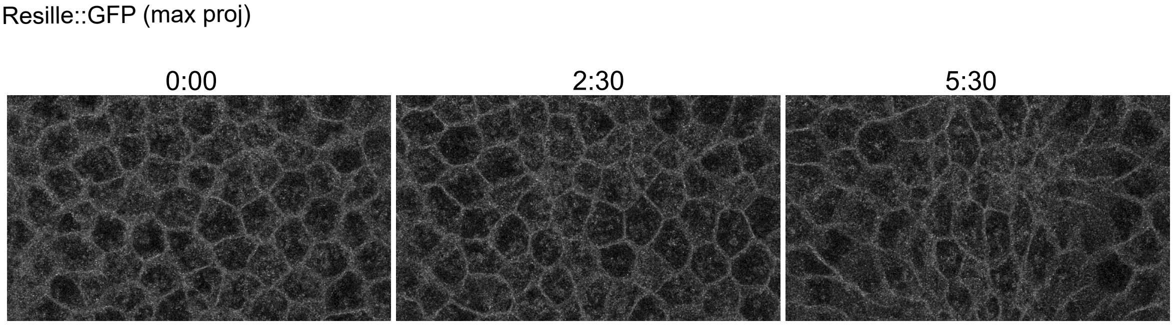


Figure S2. Resille shows uniform membrane localization during apical constriction.

Still images from time-lapse movies of Resille::GFP in the mesoderm.

**Fig. S2**

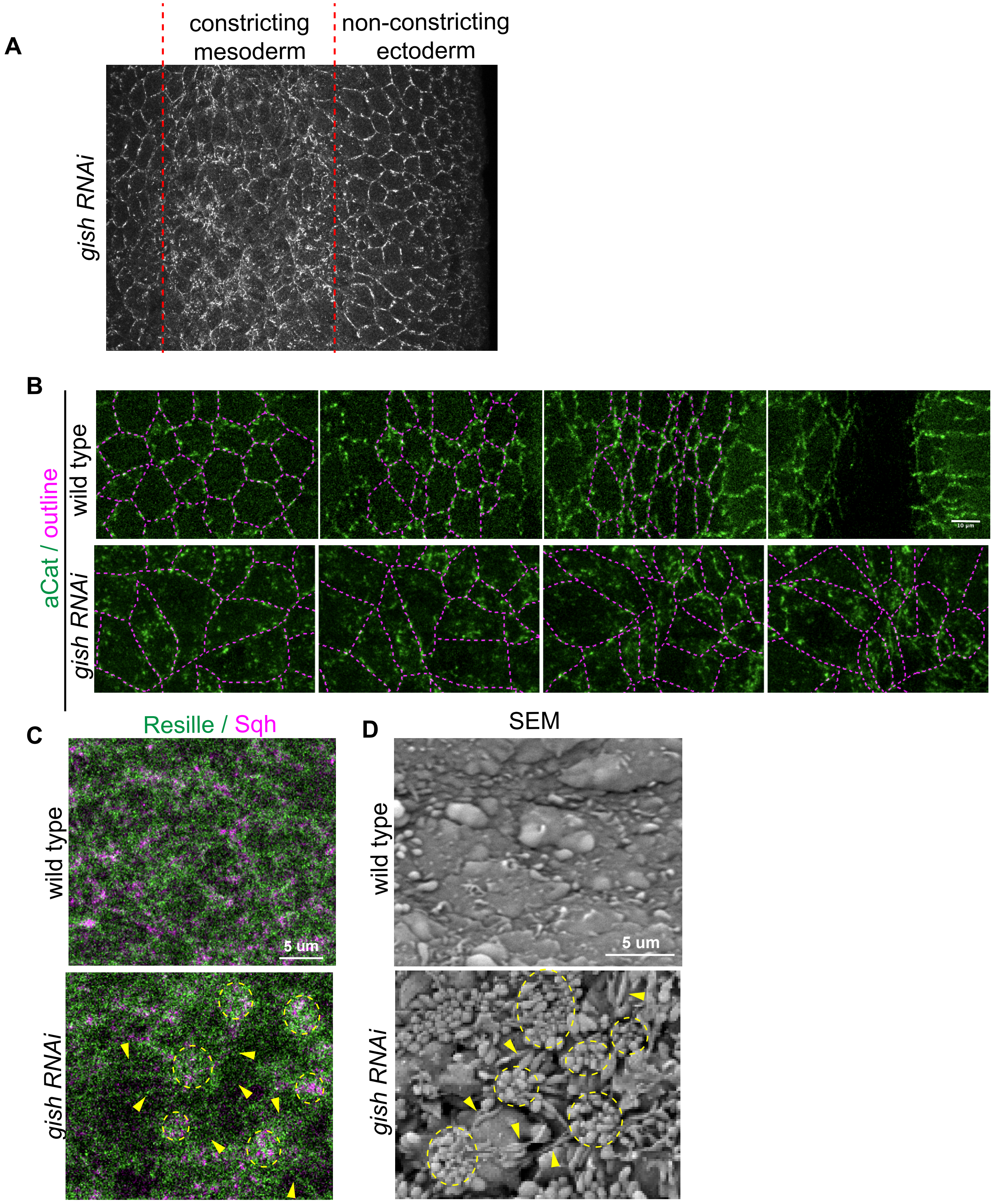
A
constricting mesoderm
non-constricting ectoderm
gish RNAi
B
aCat / outline
wild type
gish RNAi
C
Resille / Sqh
wild type
5 um
gish RNAi
D
SEM
wild type
5 um
gish RNAi

**Fig. S3**

Figure S3. Loss of Gish leads to the accumulation of adherens junctions on top of mesodermal cells during apical constriction.

(A) gish RNAi embryos immunostained for Arm. Red dotted line shows the boundary of mesoderm and ectoderm. (B) Still images from time-lapse movies of aCat::GFP in the mesoderm of wild-type and gish RNAi embryos, respectively. Dashed magenta line shows the cell boundary based on deep optical section a-Catenin signal. Scale bar: 10 μm. (C) Still images from time-lapse movies of Resille::GFP in the mesoderm of wild-type and gish RNAi embryos during apical constriction. (D) SEM images of constricting embryos of wild-type and gish RNAi embryos. Yellow dotted line: membrane blebs; Yellow arrow: gap between cells; Scale bar: 5 μm.
